# Phylogenize2: robust phylogenetic methods link genes to phenotypes across host-associated and environmental microbiomes

**DOI:** 10.64898/2026.07.15.738685

**Authors:** Kathryn Kananen, Nghi Tran, Patrick H. Bradley

## Abstract

In microbiome studies, associations between microbial functions and the environment are often confounded by phylogeny. While some methods explicitly account for this confounder, they require information about genome content, limiting their use in biomes where few genomes have been available. To make these methods more universally accessible, we have developed Phylogenize2, a redesigned phylogeny-aware tool for linking microbial gene families to abundance phenotypes. Phylogenize2 integrates large metagenome-assembled genome collections, including both biome-specific collections from MGnify and a broadly sampled general purpose database, GlobDB, to substantially expand species coverage, allowing its application in environments like the mouse gut and ocean. In addition, by default, Phylogenize2 uses a new robust phylogenetic testing framework that has been optimized for microbial abundance data, while also allowing the use of other comparative methods such as POMS. In an experimental mouse study, Phylogenize2 identifies that *Muribaculaceae* with higher abundance on a high-fat diet are enriched for proteins in the thioredoxin family, with likely roles in oxidative stress. When we apply Phylogenize2 to a polar ocean study, we find that a molybdenum-dependent PaoABC/YagTSR-like aldehyde oxidoreductase system differentiates mesopelagic from surface-dwelling *Flavobacteriaceae*, suggesting that aldehyde detoxification may be important for organisms that degrade marine snow. Together, these results show that Phylogenize2 expands phylogeny-aware microbiome analysis beyond the human gut and can provide insight into the genetic basis of microbiome-encoded traits in diverse environments.

**Importance:** Microbiome studies often set out to identify which microbes are more or less abundant across environments, but these patterns can be difficult to interpret. Phylogenize2 is an open-source software package that allows researchers to ask whether individual microbial gene families are associated with the environment across independent branches of the microbial tree of life. By incorporating large collections of genomes from uncultivated microbes, as well as modern statistical methods designed for microbial abundance data, Phylogenize2 makes this approach practical for microbiomes beyond the human gut, including in model organisms like lab mice and free-living environments like the ocean. We also provide a pipeline that allows the use of new genome collections. In two case studies, we demonstrate that Phylogenize2 effectively prioritizes specific genes and pathways from metagenomic data, thereby leading researchers from changes in microbial abundance to more biologically interpretable explanations.

## Introduction

Microbial communities both reflect and shape their environments. The sequencing revolution has made these communities legible: we can read the genomes of new taxa without cultivation, and can quantify how these taxa change across conditions, like health versus disease, or a geochemical gradient. However, an observed change in microbial abundance does not necessarily suggest a specific hypothesis for why this change happens, especially in microbes whose physiology is less well understood.

Microbial gene products, in contrast, have specific molecular functions, leading more naturally to testable hypotheses. These functions may not always be known — indeed, we still have much to learn about gene function in the microbiome^1–4^ — but this also means that gene (or gene family) associations can point us to new biology and can help prioritize which parts of this functional space to characterize. Despite these advantages, however, moving to the gene family level presents an important challenge: distinguishing genes that are directly associated from “passenger” genes that are indirectly correlated. In particular, if a clade of microbes is on average correlated with the environment, genes shared by that clade will also be correlated, even if they explain little variation within that clade or across the rest of the tree.

Phylogenetic comparative methods, like phylogenetic independent contrasts^5^ and phylogenetic regression (or phylogenetic generalized least squares)^6^, were developed to account for this type of confounding in ecological studies. Methods developed specifically for microbiome data, like Phylogenetic Organization of Metagenomic Signals (POMS)^7^ and version 1 of *phylogenize*^8^, have already shown promise for identifying microbial genes with more explanatory power than non-phylogenetic methods, which have much higher false positive rates. For example, phylogenetic methods have been used to identify the Glutamate-Dependent Acid Resistance (GAD) system as among the most enriched gene families in gut microbes^9^, and to link quinol oxidoreductases (which have a function in oxidative and nitrosative stress tolerance) to microbial abundance in obese individuals^7^.

However, in general, phylogenetic methods remain underutilized in the microbiome. One reason is statistical: metagenomic sequencing data pose unique challenges, and microbial differential abundances may not necessarily fit common models of trait evolution, motivating the development of revised methods specifically designed to be used on microbiomes (see Companion Article). Another major problem with using phylogenetic methods to map genes to microbial abundances is that they require information about which taxa encode particular gene families, meaning that they require (pan)genomes for each species. Until very recently, we have lacked genomes for the majority of microbes from many environments of interest, such as soil, the mouse gut, or even the human gut outside of certain well-studied groups^10^. Over the last several years, however, assembling genomes directly from mixed communities (yielding metagenome-assembled genomes, or MAGs) has become much more feasible. While completeness and contamination of MAGs still pose challenges for some analyses^11–13^, MAG collections have nevertheless been transformational for microbial genomics, and extensive collections now exist for host-associated^14–16^ and free-living^16^ microbiomes.

Here we present Phylogenize2, a tool that addresses the issues identified above. Phylogenize2 is a complete overhaul of the original *phylogenize*, including a new statistical test designed specifically for this application (see Companion); it is also capable of applying other phylogenetic methods, like POMS^7^, on the same data. Phylogenize2 also expands the scope of phylogeny-aware microbiome analyses by incorporating large biome-specific collections of MAGs from the continually updated MGnify Genomes^16^ resource, as well as a broader collection, GlobDB^17^, which spans the bacterial tree of life. This results in orders of magnitude more coverage of microbial species and pangenomes without requiring users to construct and annotate their own genome assemblies (although custom databases can also be added). We demonstrate that, using these collections, Phylogenize2 can be applied not only to, e.g., the human gut, but also to model organism gut microbiomes and free-living marine populations. Finally, Phylogenize2 is designed for practical use: installation is available through a Bioconda^18^ recipe, and a Snakemake^19^ workflow has been made available for users to generate their own custom databases.

## Results

### Integrating large-scale assembly collections enables “batteries-included” analyses

**Figure 1:**
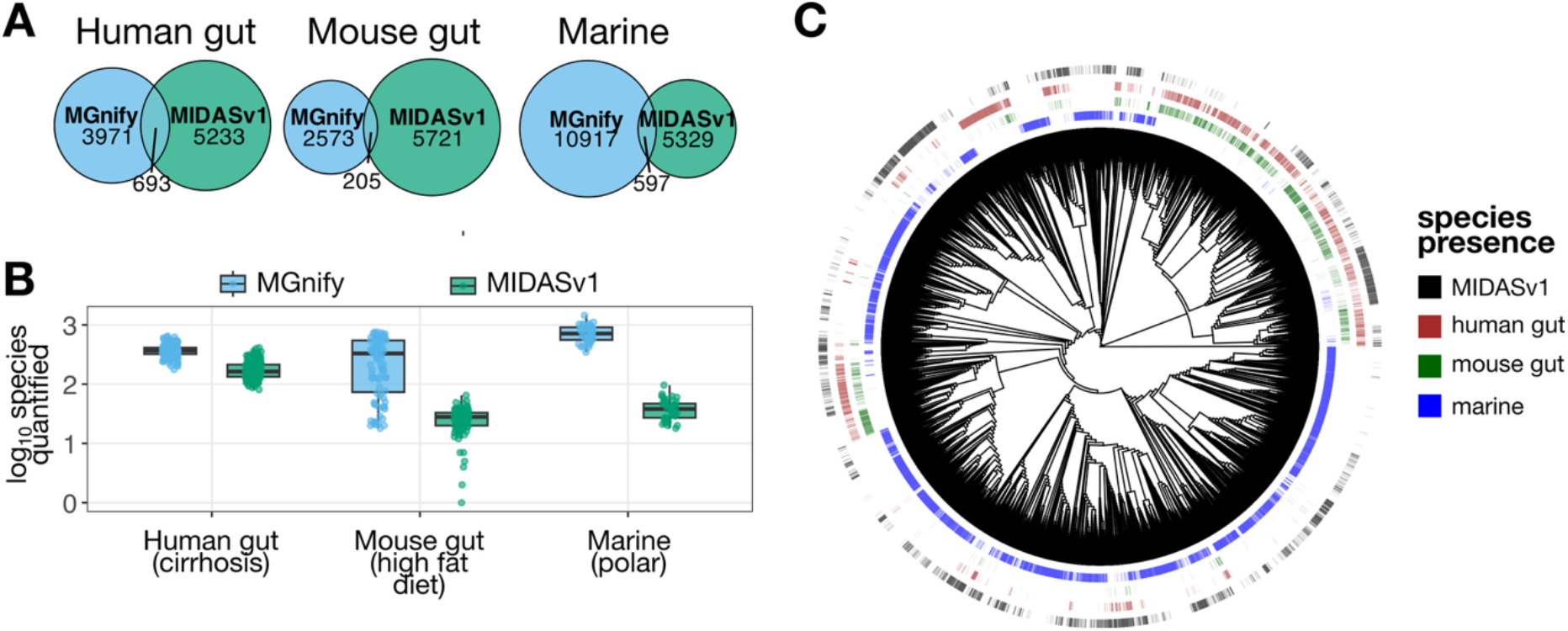
Phylogenize2 incorporates MGnify databases that sample a much larger range of taxonomic diversity. A. Overlap between species in MGnify^16^ databases specific to the human gut, mouse gut, or marine environments to MIDASv1^10^, the database used in version 1 of *phylogenize*^8^. Sourmash^20^ hashes were used to estimate ANI between pairs of genomes (see Methods). B. Log_10_ unique species detected per sample in three different public metagenomic datasets (a human case-control study of liver cirrhosis, a controlled mouse study of high-fat diet, and an observational study of polar marine samples). Samples were quantified against either the MIDASv1 database (red, right) or the appropriate MGnify database (blue, left). MIDASv1 samples were quantified using v1.2 of the tool MIDAS^10^, while the MGnify samples were quantified using Kraken2^21^ and Bracken^22^ with the provided databases. C. Phylogenetic comparison of the MIDASv1 database with three MGnify collections. The tree is derived from GlobDB^17^. Genomes were matched to tree tips using Sourmash to estimate ANI (see Methods). Only tips matching at least one database were retained for visualization.

As noted, phylogenetic methods require estimates of microbial gene content per species. In version 1 of *phylogenize*, these estimates were derived from pangenomes in the MIDASv1^10^ database, which mainly included isolate genomes from 5,926 species available through the PATRIC^23^ (now BV-BRC) database in March 2015^10^. While these genomes were high-quality, coverage was poor for many communities of interest to microbial ecologists. For example, while over 60% of reads from American human gut microbiomes could be mapped to a genome in the database, this number was less than 5% even for the mouse gut microbiome, and even worse for soil and marine environments.^10^ Fortunately, large-scale collections of metagenome-assembled genomes (MAGs)^14–16^ have resulted in expanded databases with dramatically better coverage.

Phylogenize2 builds on a collection of biome-specific databases from MGnify^16^, which includes catalogs for several body sites in both humans and model organisms, as well as environments like marine and soil. Many species have multiple genomes, allowing us to compute the frequency with which a gene family is observed in the pangenome, as opposed to just presence or absence (as in version 1 of *phylogenize*). This also helps guard against contamination of individual MAGs^11,24^. We also incorporated a large general database, GlobDB v226^17^, which contains representative genomes for 306,260 species drawn from 14 sources. In this collection, gene content is estimated from a single genome, which is less accurate; however, this database allows applications to environments not currently covered by MGnify.

We harmonized these databases by mapping protein clusters from each database to a cascading set of common references. Initially, amino acid sequences were mapped against UniRef50^25^ using MMSeqs2^26^; sequences that did not match were next mapped against UHGP-50^14^, a human gut catalog, and then FESNov^4^, a database of uncharacterized prokaryotic gene families that show evolutionary evidence of conservation. Finally, any remaining sequences were clustered *de novo* at a 50% amino acid identity level using MMSeqs2^26^ (Supplemental Figure 1). Annotations for all of these sequences were generated via HMM searches against KEGG Orthology families^27^, using the heuristic algorithm from anvi’o^28^ to improve sensitivity^29^.

We compared the coverage of these new databases to the MIDASv1 database from the original version of *phylogenize*, focusing on the human gut, mouse gut, and marine databases as examples. First, we used Sourmash^20^ to estimate pairwise average nucleotide identity (ANI) between genomes, using a species-level cutoff of ≥95.86% ANI (see Methods). Because MIDASv1 is not biome-specific, we did not expect most of its 5,926 species to be shared by any individual environment. Conversely, we found that most species in each biome-specific catalog had no match in MIDASv1. The MGnify catalogs we tested contained 6.5 times (human gut), 13 times (mouse gut), and 19 times (marine) as many species as MIDASv1 had in the same environment (**Figure 1A**).

We next determined how this increase in database coverage might translate into quantification of species from a real metagenomic dataset. We selected three test datasets — human gut cirrhosis metagenomes^30^, mouse gut metagenomes from a high-fat diet study^31^, and polar marine metagenomes^32^ — and classified them against the MIDASv1 and the MGnify databases. In all three cases, we observed substantial improvement, with 2.1, 12.2, and 18.5 times as many species quantified in the human cirrhosis, mouse microbiome, and Tara Oceans dataset, respectively (**Figure 1B**).

Finally, we assessed how this increased diversity was distributed across the bacterial tree of life. To do so, we again used Sourmash to map species from the MGnify databases and from MIDASv1 to GlobDB. This analysis showed that the species unique to the new databases were not simply close neighbors of species already covered by MIDASv1 but included entire clades that were previously underrepresented or absent (**Figure 1C**). This was especially true for the marine collection but was still the case for both mouse and even human gut. Several collections also included archaea, which were not included in MIDASv1 (Supplemental Figure 2: Phylogenetic comparison of the MIDASv1 database with three MGnify collections across archaea (phylogenetic tree from GlobDB).).

### Overview of Phylogenize2 pipeline

**Figure 2:**
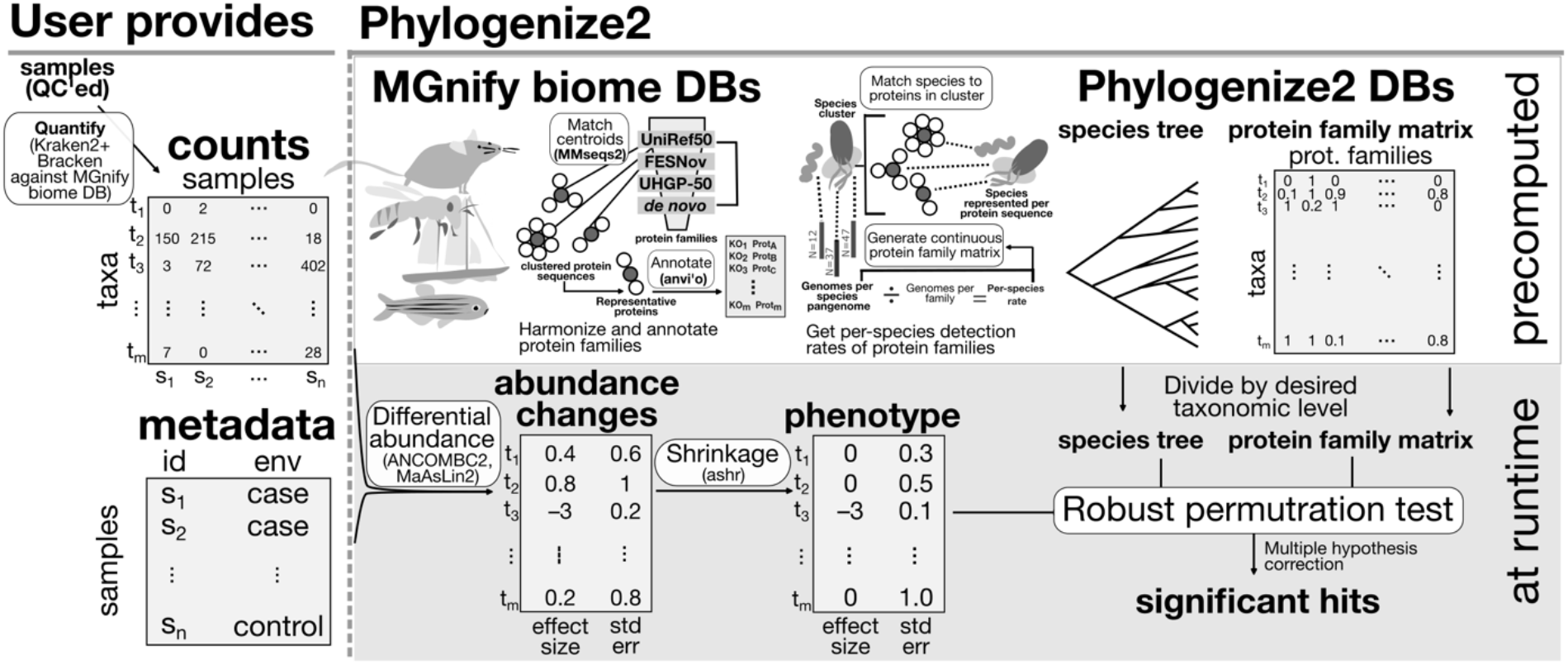
Overview of Phylogenize2. Left: the user provides a count matrix of taxa that have been quantified against one of the databases in Phylogenize2 (MGnify biome databases or GlobDB), along with a metadata table showing which samples belong to each environment. Top right: Phylogenize2 provides the databases necessary to perform the phylogenetic associations. First, the MGnify biome DBs are harmonized by matching protein sequence centroids against a common, cascading set of references, *de novo* clustering any unmatched sequences. Proteins are annotated against KEGG Orthologs using anvi’o. For each species, we quantify the frequency of detecting each protein family across individual genomes, yielding a continuous matrix with values between zero (absent) and one (core). We also provide ultrametric species trees. Bottom right: At runtime, Phylogenize2 estimates abundance changes between environments from the user’s count and metadata tables, performs adaptive shrinkage on the results, and then performs robust permutration tests to explain this phenotype in terms of protein families using the tree. Finally, p-values from these tests are adjusted for multiple testing and a report is generated.

An overview of Phylogenize2 can be found in **Figure 2**. Phylogenize2 only requires the user to provide species abundances (which can be computed using the Kraken2^21^/Bracken^22^ databases provided by MGnify^16^ or the Sylph^33^ database provided by GlobDB^17^), as well as sample metadata that show, e.g., which samples belong to which group. Phylogenize2 provides the other needed components for phylogenetic regression, namely 1) a database of pangenome content across species, and 2) a set of pre-computed phylogenetic trees. Phylogenize2 then computes differential abundances (currently, this can be done automatically via ANCOM-BC2^34^ or MaAsLin2^35^, or users may pre-compute them using any other method that returns both an effect size and a standard error) and performs adaptive shrinkage with ashr^36^. Finally, Phylogenize2 applies the “robust permutration” test (see Companion) with the aim of linking gene families from the database to differential abundance phenotypes. Phylogenize2 automatically generates a report including visualizations of the association scores, significantly associated genes, and their enrichment for KEGG metabolic pathways.

To illustrate how Phylogenize2 can be productively applied to other environments, we applied Phylogenize2 in case studies from two environments that were poorly covered by the first version, the mouse gut (comparing a high-fat or standard diet)^31^ and the polar marine environment (comparing the surface and mesopelagic zone)^32^. We also applied Phylogenize2 to a dataset that looked at the human gut in cirrhosis vs. normal liver function^30^, the results of which are described in a companion paper (see Companion). By default, Phylogenize2 uses robust permutration but also allows the user to apply other methods including POMS and an uncorrected linear model, allowing a direct comparison. On these datasets, the robust permutration method of Phylogenize2 is more sensitive than POMS while remaining much more specific than the linear model in the cirrhotic and marine datasets; interestingly, however, Phylogenize2 actually identifies more significant hits than the linear model on the mouse data (**Table 1**).

**Table 1:** Number of significant positive and negative associations found by Phylogenize2, POMS, and a linear model.

| Dataset and Taxon | Direction of Association | Phylogenize2 | POMS | Linear model |
| --- | --- | --- | --- | --- |
| **Cirrhosis**<br>(*Lachnospiraceae*) | Positive | 242 | 35 | 9391 |
|  | Negative | 124 | 0 | 4913 |
| **Marine**<br>(*Flavobacteriaceae*) | Positive | 771 | 1 | 5528 |
|  | Negative | 379 | 9 | 4602 |
| **Mouse**<br>(*Muribaculaceae*) | Positive | 52 | 0 | 21 |
|  | Negative | 27 | 0 | 3 |

**Figure 3:**
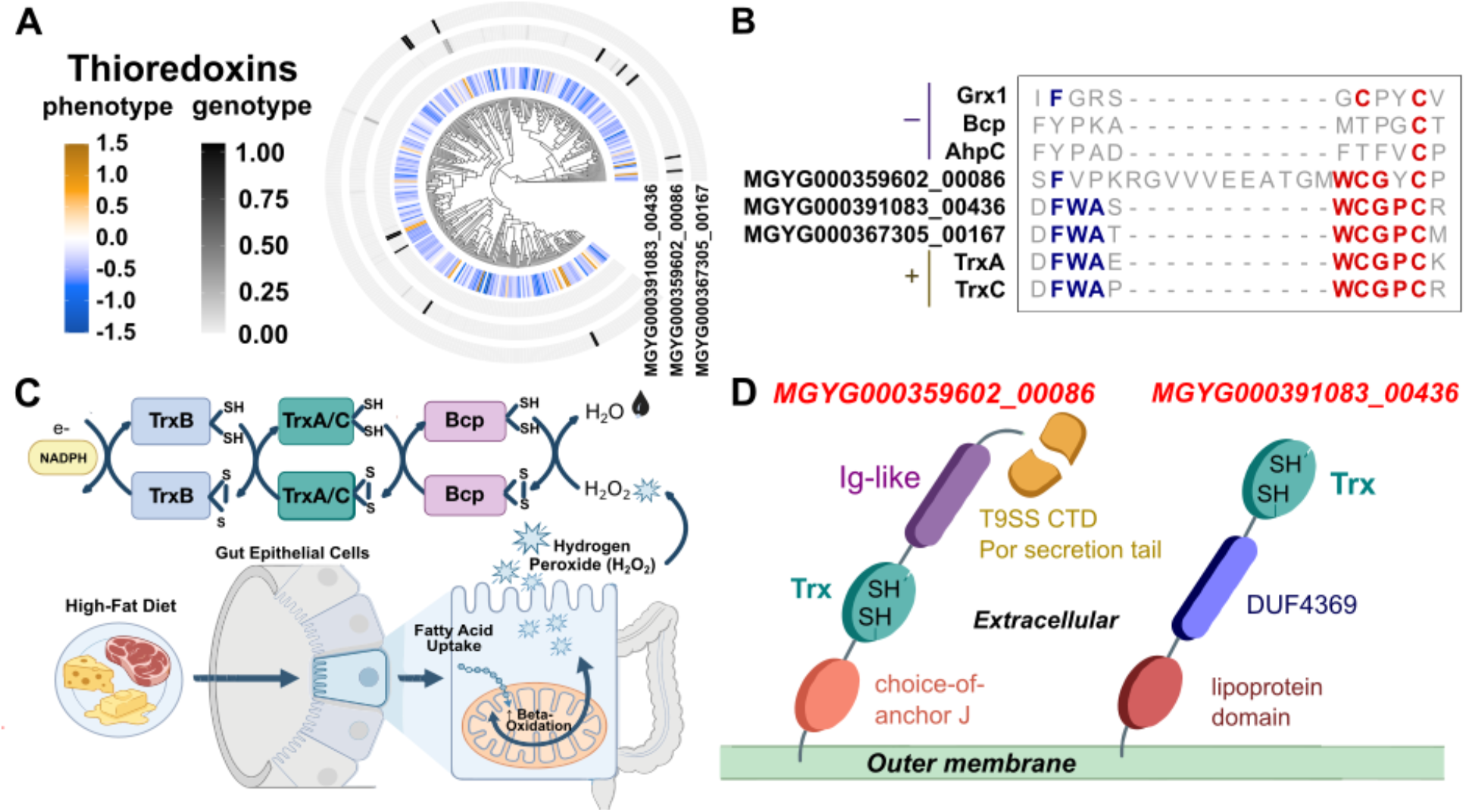
Identification of different thioredoxins linked to *Muribaculaceae* abundance in mouse models of high-fat diet (HFD). A) Differential abundance phenotype (blue to red) plotted against the genotype (white to black) for three significant hits from Phylogenize2 on the *Muribaculaceae* tree. B) Multiple alignment results highlighting conserved sites and canonical active sites of three target hits and other thioredoxin superfamily members in *E. coli*. Grx1: Glutaredoxin_1, Bcp: Peroxiredoxin, AhpC: Alkyl hydroperoxide reductase subunit C, TrxA: Thioredoxin_1, TrxC: Thioredoxin_2. C) Hypothetical reaction mechanism of the canonical thioredoxin 1/2 (TrxA/C) (MGYG000367305_00167 hit) localized in cytoplasm, along with other thioredoxin-containing partners, TrxB: Thioredoxin reductase, Bcp: Peroxiredoxin, showing detoxification of peroxide and electrons being passed to maintain all members in a reduced, active form. D) InterPro domains of MGYG000359602 and MGYG000391083 showing localization and tethering at the membrane to perform the roles shown in panel C. Ig-like: Immunoglobulin-like, DUF4369: domain of unknown function 4369. Panels C and D were created using FigureLabs (FigureLabsAI, https://www.figurelabs.ai).

### Case study: In mice, thioredoxins may protect *Muribaculaceae* against host-derived peroxides released on a high-fat diet

Phylogenize2 was applied to mouse gut microbiome samples from a high-fat diet (HFD) experimental study^37^ using the MGnify mouse gut database. We focused on the *Muribaculaceae* family, which is known to confer beneficial effects on host health yet which remains less well studied than its closest relatives, the *Bacteroidaceae*. Consistent with previous reports showing that *Muribaculaceae* have reduced abundance in mouse models of HFD^38^, our analysis revealed a predominantly decreasing trend in *Muribaculaceae* taxonomic abundance (**Figure 3A**). Using Phylogenize2, we identified 52 gene families that were significantly positively associated (q < 0.05) (Supplemental Table 1). POMS did not recover any associations, and interestingly, the linear model returned only 21 associations, 9 of which overlapped with Phylogenize2’s.

Among the top significant hits, three gene families were identified as being positively associated with HFD conditions and appeared to all contain thioredoxin domains: these were UniRef50_A0A4Z0V8U3 (MGYG000391083_00436 in the original MGnify database), UniRef50_A0A2M8N864 (MGYG000367305_00167), and UniRef50_A0A7J0ABB5 (MGYG000359602_00086). These proteins were nominally significant (uncorrected p < 0.05) using the linear model, but only one was significant after correction (UniRef50_A0A7J0ABB5). Thioredoxins typically play roles in the oxidative stress response; notably, high-fat diets have also been found to increase oxidative stress in the gut, driven by enhanced fatty acid oxidation, impaired epithelial mitochondrial function, and the release of oxygen and reactive oxygen species (ROS) into the normally anaerobic gut^39–41^.

As some of these UniRef50 sequences appeared to be fragments, we used the full-length sequences from the original MGnify database (MGYG IDs) to construct alignments with thioredoxin superfamily members. This demonstrated that the targets shared the highest similarity with thioredoxin 1 and thioredoxin 2 (TrxA and TrxC), particularly within the canonical WCxxC active-site motif, compared with other members of this family potentially involved in H_2_O_2_ detoxification (**Figure 3B**).

Although all three targets were identified as part of the thioredoxin superfamily, InterPro^42^ functional domain analysis and BLAST^43^ annotation of their genomic neighborhoods suggested distinct subcellular localization patterns and functional roles. MGYG000359602_00086 and MGYG000391083_00436 were predicted to be exported to the outer membrane through two different secretion mechanisms, whereas MGYG000367305 was predicted to remain cytoplasmic. Unlike the membrane-associated targets, MGYG000367305_00167 had no signal peptide sequence, and its neighborhood contained genes annotated as having functions in DNA repair (Supplemental Table 8). This protein therefore likely functions as a canonical thioredoxin that may be induced to prevent DNA damage by peroxides. Specifically, the MGYG000367305_00167 could maintain the peroxiredoxin Bcp^44^ in its reduced, active state through electron transfer from these targets, enabling the conversion of H_2_O_2_ into water (**Figure 3C**)^45^.

In contrast, the genome neighborhoods of MGYG000359602_00086 and MGYG000391083_00436 suggest a role in nutrient uptake, especially iron. Specifically, MGYG000359602_00086 was identified as a likely T9SS-secreted surface protein containing a thioredoxin domain and an Ig-like fold, attached to the outer membrane through the T9SS C-terminal domain (CTD) anchored mechanism (**Figure 3D**, left). Based on its domain similarity to the hemin-binding protein HBP35 from *Porphyromonas gingivalis,* which is also secreted via the T9SS^46^ and contains a thioredoxin^47^ domain and an Ig-like fold, as well as similarities in the genome neighborhood (Supplemental Table 6), we hypothesized that this protein may both protect surface-exposed proteins from oxidative damage while facilitating the transport of potentially reactive iron.

Similarly, MGYG000391083_00436 was also predicted to encode an outer membrane lipoprotein, but one exported through the Sec secretion pathway (**Figure 3D**, right). We also found TonB-dependent transporters in the genomic neighborhood of MGYG000391083_00436 (Supplemental Table 6), suggesting that this protein also plays a role in substrate transport. Furthermore, a recent study of three diverse Bacteroidota under iron deprivation (*B. thetaiotaomicron, P. intermedia,* and *P. gingivalis*) showed that thioredoxin proteins and TonB-dependent transporters were upregulated in 2/3 and a protein with a DUF4369 domain was upregulated in 3/3.^48^

Together, the identification of multiple thioredoxin-associated mechanisms by Phylogenize2 suggests that oxidative stress is elevated under high-fat diet conditions, driving *Muribaculaceae* to employ diverse strategies for ROS detoxification. These results also point to a potential specific role in iron acquisition, which may be connected to ROS exposure because iron in the presence of either oxygen or peroxide can drive the toxic radical-generating Fenton and Haber-Weiss reactions^49^.

### Case study: In the polar ocean, molybdenum-dependent aldehyde oxidoreductases differentiate mesopelagic from surface *Flavobacteriaceae*

**Figure 4.**
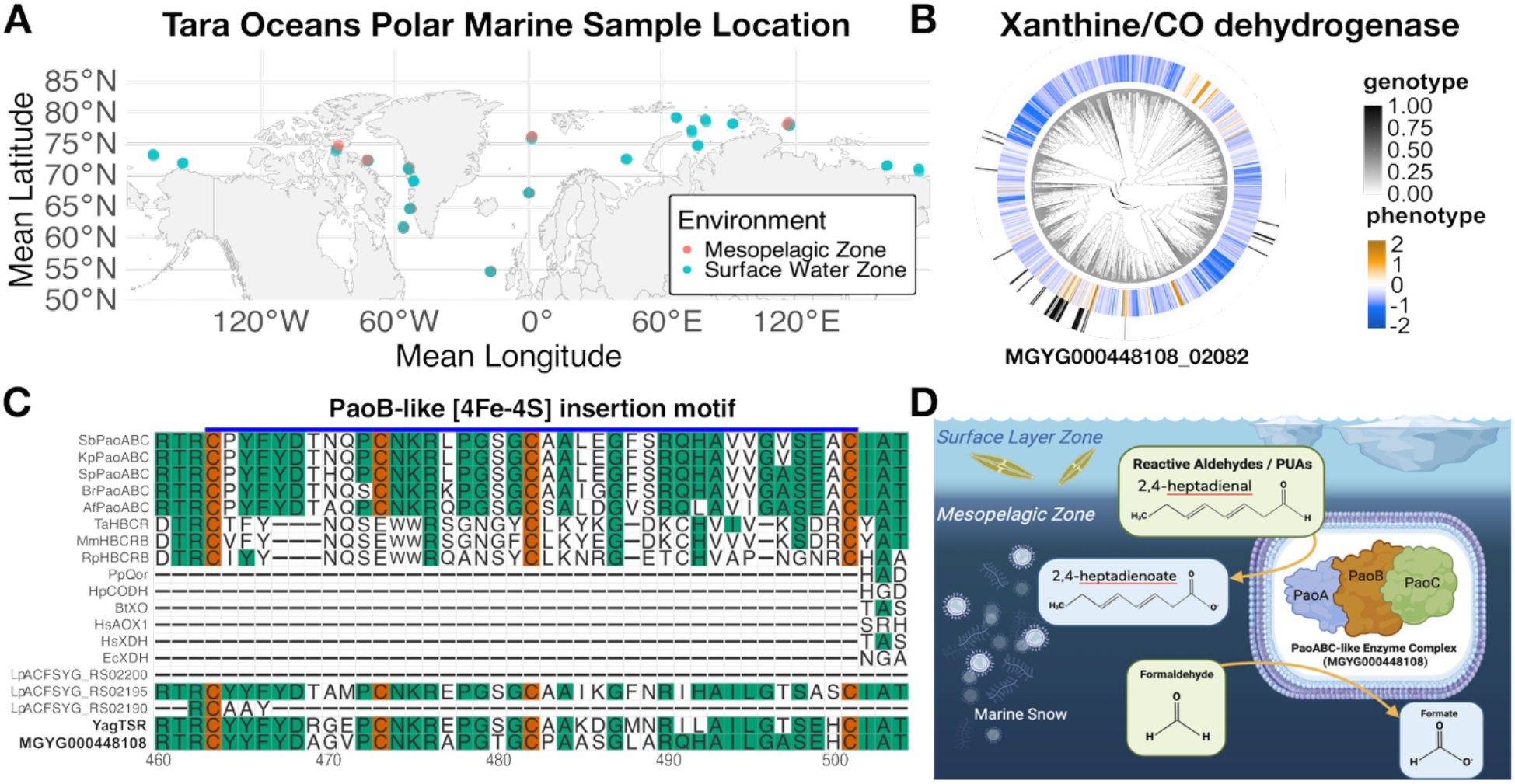
Identification of a molybdenum-dependent PaoABC/YagTSR-like system in polar marine mesopelagic *Flavobacteriaceae*. A. Mean latitude and longitude coordinates of Tara Oceans samples in polar surface water and mesopelagic collection sites. B. Phenotype (red: higher in mesopelagic; blue: higher in surface) plotted against presence (black) or absence (white) of the protein family MGYG000448108_02082. C. Multiple sequence alignment of experimentally validated members of related molybdenum-flavin enzyme families, including the aldehyde oxidoreductase FAD subunit PaoABC, 4-hydroxybenzoyl-CoA reductase B subunit HbrC, quinoline 2-oxidoreductase C subunit QorC, carbon monoxide dehydrogenase CODH, xanthine oxidase B subunit XO, aldehyde oxidase AOX1, and xanthine dehydrogenase XDH. Abbreviations indicate source organisms: Sb, Shigella boydii; Kp, Klebsiella pneumoniae; Sp, Sodalis praecaptivus; Br, Bradyrhizobium retamae; Af, Asanoa ferruginea; Ta, Thauera aromatica; Mm, Magnetospirillum magneticum; Rp, Rhodopseudomonas palustris; Pp, Pseudomonas putida; Hp, Hydrogenophaga pseudoflava; Bt, Bos taurus; Hs, Homo sapiens; Ec, Escherichia coli; and Lp, Leeuwenhoekiella polynyae. The alignment also includes the L. polynyae sequence with the highest sequence identity to the YagTSR proteins, significantly enriched YagTSR proteins, and the MGYG000448108_02082 PaoABC-like protein. Sites conserved in canonical PaoABC proteins are highlighted, including four conserved cysteine residues in the iron– sulfur-binding subunit that are retained in both YagTSR and MGYG000448108_02082. D. Hypothetical function of MGYG000448108_02082 in polar marine mesopelagic *Flavobacteriaceae* using marine snow PUAs and formaldehyde for particle degradation and detoxification. The PaoABC complex is represented as a cytoplasmic molybdenum-dependent aldehyde-oxidizing system whose precise subcellular location remains unresolved. Figure 4D was created using BioRender (https://biorender.com/d0ko35h).

To demonstrate Phylogenize2’s applicability to free-living environments, we applied it to data collected from a survey of polar marine microbiomes from the Tara Oceans project. Because metabolic lifestyle is heavily structured by depth (because of factors like oxygen, light availability, and which nutrients are available), we contrasted bacterial communities at the surface with those in the deeper mesopelagic zone.

Here, we focused on the *Flavobacteriaceae*, an abundant Gram-negative taxon in the Bacteroidota. In the mesopelagic zone (ENVO:00000213), both non-sinking and fast-sinking marine particles are sites of particulate organic carbon transformation^50^. Prior reports have shown that Flavobacteria are significantly enriched in abundance on bulk particle samples compared to small-size-fraction seawater samples in the North Pacific Subtropical Gyre^51^. In the upper mesopelagic zone in the Porcupine Abyssal Plain, the *Flavobacteriaceae* family was shown to be the second most relatively abundant family on fast-sinking particles at 70 meters^50^. Furthermore, these taxa are consistent with reported early particle colonizers that occupy a complex-polysaccharide-degrading role potentially breaking particles into dissolved organic matter^52,53^. As particles are degraded, other cellular debris and dissolved metabolites can be released from organic matter, providing new substrates for metabolism.

We found that when comparing the polar marine mesopelagic and polar surface water zones (**Figure 4A**), mesopelagic *Flavobacteriaceae* were enriched for a protein (MGYG000448108_02082) annotated as a subunit of xanthine dehydrogenase (XDH) (**Figure 4B**). However, we noted that in addition to XDH, this family includes related proteins such as carbon monoxide dehydrogenases and aldehyde oxidoreductases, which can be difficult to distinguish^54^. We therefore generated alignments of MGYG000448108_02082 with experimentally verified sequences from different proteins in the family (Supplemental File 2). This revealed substantial protein identity (max of 53.38%, min of 47.72%) across five sequences from the PaoABC complex, which is a periplasmic aldehyde oxidoreductase (**Figure 4C**; Supplemental File 2). In particular, we saw that MGYG000448108_02082 contained an additional [4Fe–4S] cluster binding motif with four conserved cysteines in the FAD-binding subunit, which is characteristic of the aldehyde oxidoreductase PaoABC complex^54^. Interestingly, MGYG000448108_02082 appears to be a potential fusion of these subunits; XDH can also exist as multiple ORFs or as a fusion. In addition, proteins annotated as YagR (K11177), YagS (K11178), and YagT (K13483) were significantly associated with mesopelagic *Flavobacteriaceae;* these also aligned to PaoABC with a high degree of homology (∼60%), as expected since these are the original names of the periplasmic aldehyde oxidase complex in *E. coli*^55^. Finally, we also saw a significant association with MobA (K03752), which is involved in the synthesis of molybdenum cofactor (MoCo); notably, PaoABC is a MoCo-dependent enzyme.

Together, these results indicate that polar mesopelagic *Flavobacteriaceae* are enriched for a molybdenum-dependent PaoABC/YagTSR-like system, which is likely involved in the oxidation of aldehyde substrates rather than carbon monoxide or purines^54^. (Note that while key residues were the most similar to PaoABC, one difference is that we did not find evidence of a Tat signal peptide, so the subcellular localization of this complex may not be periplasmic in *Flavobacteriaceae*.)

One functional explanation is that this system is linked to the degradation of organic matter particles^56^. Particles in “marine snow,” organic matter that descends from the surface to the pelagic ocean, are known hot spots of microbial activity and remineralization. Diatoms, in particular, are known to be major components of marine particles degraded by Flavobacteria^52^, and are enriched in polar marine environments^57^. In response to stress or predation, many diatoms also produce potentially toxic polyunsaturated aldehydes (PUAs) such as heptadienal, which could affect particle-attached bacterial metabolism^56^. Therefore, we would expect these particles to potentially contain phytoplankton-derived reactive aldehyde compounds (**Figure 4D**). In addition, algal-polysaccharide metabolism by marine Flavobacteria can itself generate toxic aldehyde byproducts such as formaldehyde, as shown during porphyran utilization in *Zobellia galactanivorans*, where formaldehyde detoxification is coupled to algal-polysaccharide degradation^58^. While the precise substrate of this enzyme remains to be determined, this supports the broader idea that the capacity to detoxify aldehydes and/or use them for redox chemistry may be beneficial during degradation of organic matter in the mesopelagic zone (**Figure 4D**).

When we compared against the linear model, we found that it did also call the aldehyde oxidoreductase proteins significant, but not MobA, the molybdenum cofactor biosynthetic protein; furthermore, that model yielded over 5.5K total positive associations, consistent with the high false-positive rate that has been demonstrated for uncorrected linear associations^8,9^. Conversely, POMS only found one association at q < 0.05, which did not overlap with Phylogenize2. This association would therefore have been difficult to discover using previous methods.

### Conclusions

Phylogenize2 represents several advances in applying phylogenetic models to microbiome data. First, Phylogenize2 implements state-of-the-art phylogenetic comparative methods that are designed for the microbiome, including robust permutration, which we showed to be more sensitive and specific on metagenomic abundance data than competing methods (see Companion). Next, integrating biome-specific collections that include metagenome-assembled genomes (MAGs) makes Phylogenize2 much more generally applicable than the first version of Phylogenize. Indeed, on mouse and marine data, we found that these new collections allowed us to quantify an order of magnitude more species; since each species is a single observation in phylogenetic regression, this is a major improvement in the signal we can extract from these data. Phylogenize2 can therefore be used across diverse hosts as well as in free-living environments, where much of the diversity remains uncultured. Finally, Phylogenize2 is available as a recipe in Bioconda, making it much easier to install.

We demonstrate that in real-world datasets, Phylogenize2 can be used to generate new molecular hypotheses about how microbial genes are linked to community-level phenotypes. For instance, when we applied Phylogenize2 to a study of high-fat diet in the mouse gut, we found that thioredoxin proteins, both cytosolic and at the cell surface, were strongly associated with *Muribaculaceae* abundance. Interestingly, we also saw that protein families involved in oxidative stress resistance were associated with *Lachnospiraceae* abundance in cirrhosis (see Companion). This is consistent with experimental results showing that one of the major underlying variables determining the mammalian gut microbiome composition is inflammation, a complex environmental change in which the epithelial barrier function is worsened, oxygen tension increases, and reactive oxygen and nitrogen species are formed^59,60^. We note that when applied to a large human case-control study of obesity, POMS also identified an enzyme family linked to oxidative stress tolerance (cytochrome bd oxidase)^7^.

Meta-analyses have been conducted to identify whether reproducible microbiome signatures exist, either disease-specific or pan-diagnostic^61–65^. Because individual studies are drawn from populations that can have different gut community composition at baseline (for example, because of geography^62^), methods that work at the gene level should show improved agreement across studies. However, previous meta-analyses have not used phylogenetic correction, and indeed, a meta-analysis of gut metagenomes found that the genes most strongly associated with disease were primarily markers for a small number of strains or species^65^. It is therefore interesting that we see some signs that phylogenetic methods may help us converge on a common latent environmental variable in the gut, i.e., inflammation. An important future direction will therefore be to test whether cross-disease gene associations identified using phylogenetic methods will primarily reflect this common influence of inflammation, or whether we can also identify microbial genes linked to more specific disease processes.

In addition, we showed that Phylogenize2 can be productively applied in free-living environments. In the polar ocean, our results indicated that a PaoABC/YagTSR like oxidoreductase system differentiated mesopelagic from surface *Flavobacteriaceae*. This association suggests that aldehyde oxidation may be an important trait for Flavobacteriaceae inhabiting deeper particle-associated environments where the degradation of phytoplankton derived organic matter can release reactive aldehydes and other reduced carbon compounds. Although the precise substrate and subcellular localization of the system remain unresolved, this illustrates how Phylogenize2 can help prioritize candidate metabolic mechanisms for microbial adaptations in complex environmental microbiomes.

Users of Phylogenize2 should still be aware of certain remaining limitations. Although MAG collections greatly expand environments that can be utilized by Phylogenize2, MAGs vary in completeness and contamination, which can introduce errors into estimates of gene content^11,13,66–68^. In within-species analyses, even low levels of MAG-derived pangenome contamination can cause disproportionate numbers of false positives^11^. We expect that across-species analyses, like Phylogenize2, should be more robust, as the same contaminants would have to be introduced into multiple species to bias the results. Our database pipeline also helps mitigate these issues by estimating gene-family detection rates across all genomes for a species; many contaminants are only seen in a single MAG^24^. Species represented by only a few genomes, however, remain more vulnerable to errors, and low-abundance or difficult-to-assemble taxa may still be underrepresented in the current genome collections. Analyses that use GlobDB may also be more susceptible than those that use the MGnify collections, since in GlobDB each species is represented by a single genome.

Other important caveats arise from the way protein families are defined. In both MGnify and GlobDB, protein families are constructed using the MMseqs2 linear clustering algorithm (“linclust”), which has been shown to overpartition protein families^69,70^, potentially leading to a failure to detect genes in certain taxa that are actually conserved. True all-against-all alignments yield protein families with more accurate phylogenetic profiles, which have successfully been used to identify human gut colonization factors^71^; even with improvements to compute time, however, this approach remains prohibitive to apply at the scale of multiple biomes. New, scalable tools for protein family inference are therefore needed.

Finally, as we observed with both case studies, existing protein family annotations can be inadequate (failing to differentiate the thioredoxin-containing proteins) or misleading (annotating a likely aldehyde oxidoreductase as a xanthine dehydrogenase). Here, we used manual analyses of genome neighborhoods and protein sequences to zero in on function; large-scale function prediction tools that incorporate these data sources, such as FUGAsseM^1^, are likely to improve annotation quality and save researchers time. As MAG recovery, genome curation, and genome annotation continue to improve, the Phylogenize2 databases can be updated to reduce these sources of uncertainty and expand phylogeny-aware functional analysis to additional microbial environments.

## Methods

### Metagenomic data

Metagenomic datasets representing the mouse gut under a normal or high-fat diet (PRJEB52043)^31^ and the polar ocean at varying depths (PRJEB9740) were downloaded from the European Nucleotide Archive^72,73^ and processed as described in the companion article (see Companion).

### Database processing

Phylogenize2 utilizes GlobDB v226 and 17 MGnify databases. For each database, processing and annotation were run via a Snakemake workflow (Supplemental Figure 1). All processed and annotated databases are listed in Supplemental Table 4. The workflow has been made available on GitHub for use with MGnify databases, GlobDB, or any database that follows MGnify’s internal naming conventions.

We started with protein sequences that had been clustered at the 50% amino acid identity level with the MMseqs2 “linclust” linear-time clustering algorithm^26^. We then used MMSeqs2 v17.b804f to search each protein cluster’s centroid against UniRef50, UHGP, and FESNov. For GlobDB, singleton protein clusters (i.e., with only one member) were added back, as these were removed by default. MMseqs2 was run iteratively, raising the sensitivity from 3 to 7 to optimize clusters mapped versus runtime (Supplemental Figure 3). For all databases, any clusters that were not mapped to one of the three reference databases were clustered together using MMseqs2’s “linclust.”

All protein clusters for each database were then annotated using anvi’o. For the MGnify pangenomes, a modified anvi’o fork (commit 0933fd1) was used, which allowed for amino acids to be annotated directly without starting from nucleotide sequences. The KEGG KOfam database was downloaded on 2025-02-04. For GlobDB, pre-existing KO annotations from anvi’o were used.

The phylum-level trees from each database were made ultrametric by calculating relative evolutionary divergence^74^ as implemented in the R package “castor”^75^, and were otherwise unmodified. Protein family detection rates were calculated for each MGnify database, yielding a matrix with values ranging from 0 (not detected) to 1 (always detected). For a given 50% ID protein cluster and a given species, the detection rate was defined as the number of genomes encoding proteins assigned to that cluster divided by the total number of genomes for that species. Since GlobDB was dereplicated at the species level, a binary matrix (0: absent, 1: present) was generated instead for that database.

### Database comparisons

Sourmash^20^ v4.9.4 was used to estimate average nucleotide identity (ANI) between pairs of genomes. First, to determine an appropriate estimated ANI threshold to distinguish taxa at the species, genus, and family levels, we compared the MGnify mouse gut, marine, and human gut databases to themselves. For this comparison, the marine and mouse gut databases were subsampled by taking the top 50 most complete genomes per representative species. The human gut database was subsampled by taking the top 10 most complete genomes per representative species. Next, k-mer hash databases were generated, using a k-mer length of 31 as recommended. We found that 90% of genomes from the same species were above an ANI cutoff of 95.86% (Supplemental Figure 4; Supplemental Table 5). Next, k-mer hash databases (again using 31-mers) were generated for each species representative in MIDASv1, GlobDB v226, and the MGnify genome collections. Each database was compared to MIDASv1 (**Figure 1A**) and GlobDB (**Figure 1B**), counting the best ANI score above 95.86% as a match. For each of our three example datasets (cirrhosis, high-fat diet, and polar marine metagenomes), we also computed the number of unique species that could be quantified in each sample, either using Kraken2 and Bracken and the appropriate MGnify database, or using MIDASv1 as in version 1 of *phylogenize* (**Figure 1C**).

### Case studies

#### Prioritizing and visualizing gene hits

For the mouse and marine datasets, we examined all significant (q < 0.05) positive associations and prioritized the 50 with the broadest phylogenetic distribution (Supplemental Table 6: Faith’s phylogenetic diversity (PD) of top mouse Muribaculaceae hits., Supplemental Table 7: Faith’s phylogenetic diversity (PD) of top marine Flavobacteriaceae hits.). We quantified this using Faith’s phylogenetic diversity^76^ (“picante”^77^ R package) applied to the taxa where the gene was present or absent (whichever was rarer). In addition to the KEGG Orthology annotations above, more functional annotations were retrieved by searching for the cluster representative identifier in UniProtKB and UniParc. For clusters of interest that were annotated as “unknown” or “uncharacterized,” the corresponding FASTA sequences were extracted and queried against the NCBI nr^78^ database using BLASTP^43^, with the top five relevant matches retained. To visualize the relationship between gene presence-absence patterns and the trends in phenotype (i.e., log differential abundance), both were displayed as heatmaps around the phylogenetic tree using the ggtree^79^ and ggtreeExtra^80^ R packages.

#### Mouse gut in high-fat diet: thioredoxins

We retrieved the amino acid sequences from the mouse gut MGnify database^16^ of the three *Muribaculaceae* thioredoxin-containing proteins that were significantly associated with high-fat diets. Sequences were submitted to InterPro^42^ to predict domain architecture and potential localization. Protein localizations were further investigated using DeepTMHMM^81^ to predict if each protein was secreted (MGYG000391083_00436 and MGYG000359602_00086), membrane-associated, or intracellular (MGYG000367305_00167).

Because these hits could be confidently assigned to the thioredoxin family but not to a specific homolog, we performed multiple sequence alignments against representative members of the thioredoxin family with diverse roles in redox homeostasis, including thioredoxin, thioredoxin reductase, alkyl hydroperoxide reductase subunit C, glutaredoxin, and peroxiredoxin. Alignments were examined for a conserved CxxC active-site motif shown to be critical for thioredoxin family proteins^82^.

After confirming that all hits were thioredoxin homologs, with the highest degree of conservation with TrxA and TrxB, we examined the genomic neighborhood of each hit by retrieving flanking sequences from the MGnify genome where the protein family centroid was encoded. We combined any existing annotations with BLAST searches of neighboring genes (against UniProt^83^ and NCBI BLAST^43^), with particular attention to those located on the same strand.

#### Marine mesopelagic layer: xanthine/CO/aldehyde dehydrogenases

To gain more insight into the likely function of the putative xanthine/carbon monoxide dehydrogenase subunit and YagTSRs that Phylogenize2 identified, we aligned them using MAFFT v7.526^84^ against 14 previously experimentally verified sequences in the same superfamily, comprising xanthine dehydrogenases, carbon monoxide dehydrogenases, and aldehyde oxidases, and visualized the results using ggmsa v1.6.0^85^ in R. Previously, PaoABC proteins were shown to have a 4Fe–4S cluster not found in CO or xanthine oxidoreductases; we therefore visualized the 4Fe–4S cluster-binding motif, showing that it was present in both the YagTSR complex and the MGYG000448108_02082 protein that Phylogenize2 identified. The full alignment, showing that MGYG000448108_02082 aligns to the entire YagTSR complex as opposed to a single subunit, can be found in Supplemental Figure 5.

## Availability

Phylogenize2 is available through GitHub (https://github.com/pbradleylab/phylogenize) and Bioconda. The Phylogenize2 database processing pipeline is available at https://github.com/pbradleylab/phylogenize-db-prep. Processed Phylogenize2 databases can be downloaded from the Phylogenize community page on Zenodo (https://zenodo.org/communities/bradley_phylogenize/records?q=&l=list&p=1&s=10&sort=newest). Scripts for figure generation are available at https://github.com/pbradleylab/phylogenize_v2_2025_paper. The modified version of anvi’o for protein annotation that we used is available at https://github.com/Kekananen/anvio.

## Acknowledgements

We wish to acknowledge other members of the Bradley lab for helpful discussions. Funding for this study was provided by The Ohio State University (startup funds to P.H.B.) and by the National Institutes of Health (grant R35GM151155 to PHB). The Ohio Supercomputer Center provided high-performance compute resources.

## Notes

https://zenodo.org/communities/bradley_phylogenize/records?q=&l=list&p=1&s=10&sort=newest

### Competing Interest Statement

The authors have declared no competing interest.

