## Supplementary figures and images for "Phylogenize2: robust phylogenetic methods link genes to phenotypes across host-associated and environmental microbiomes"

### Supplemental Figure 1

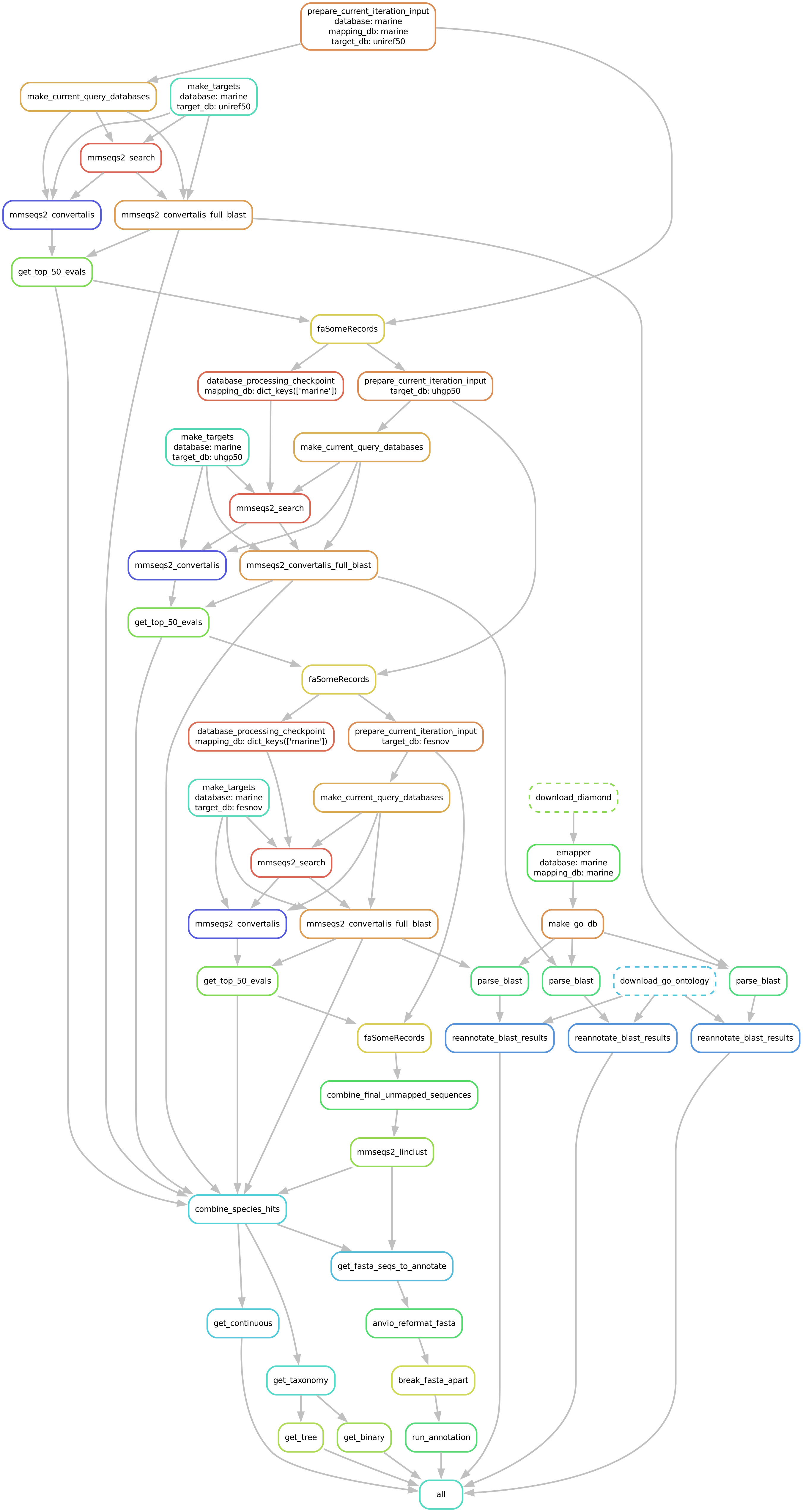

### Supplemental Figure 2

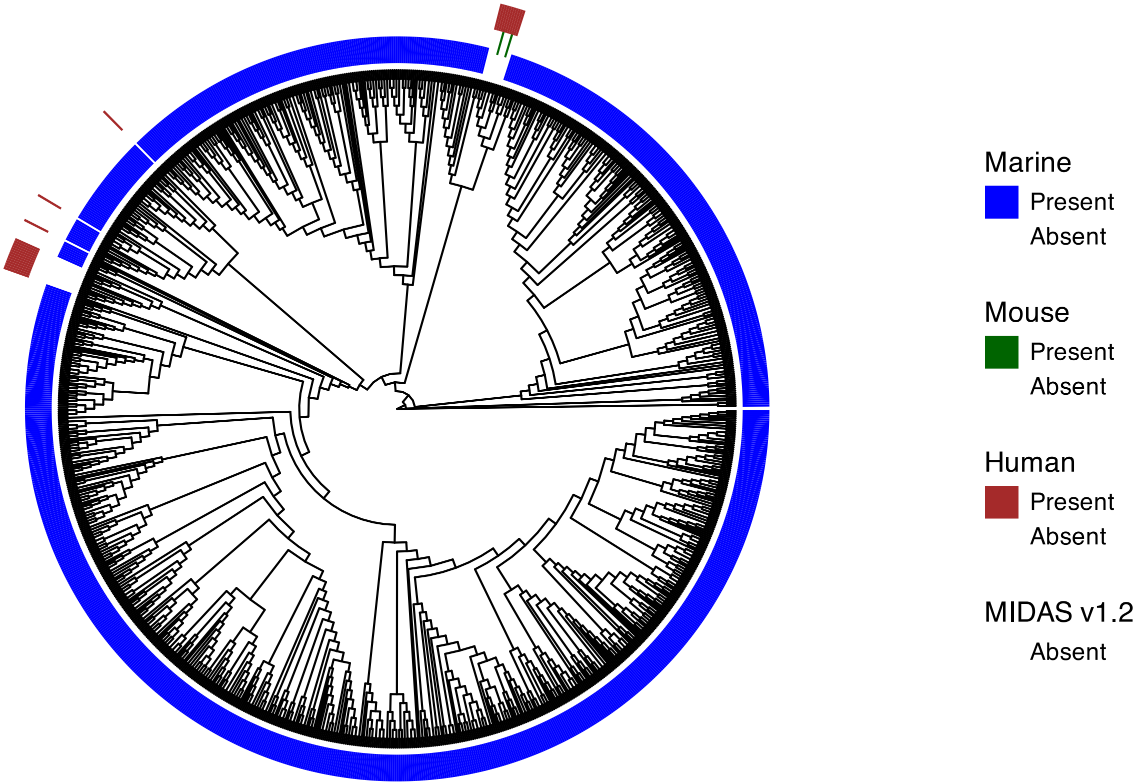

### Supplemental Figure 3

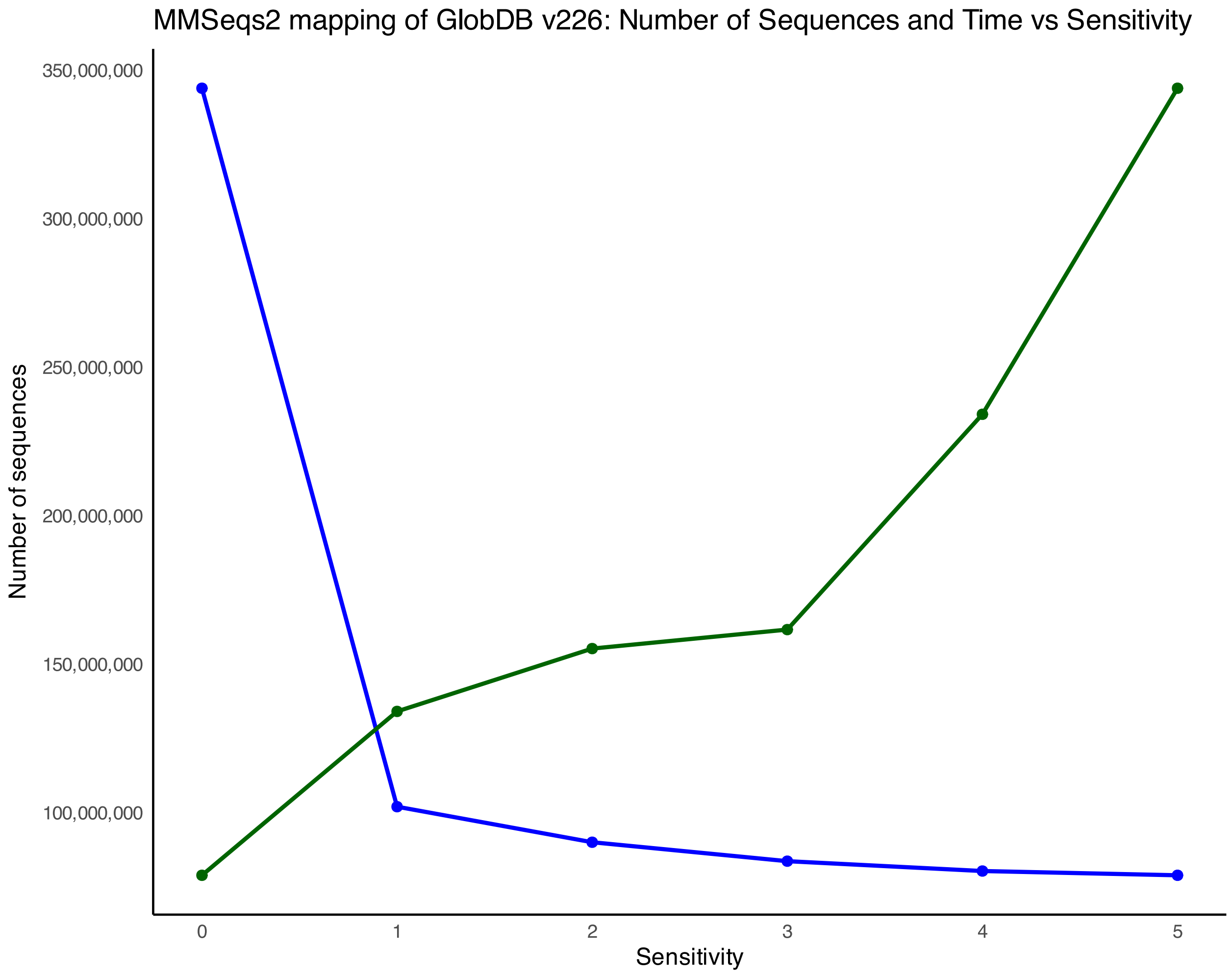

### Supplemental Figure 4

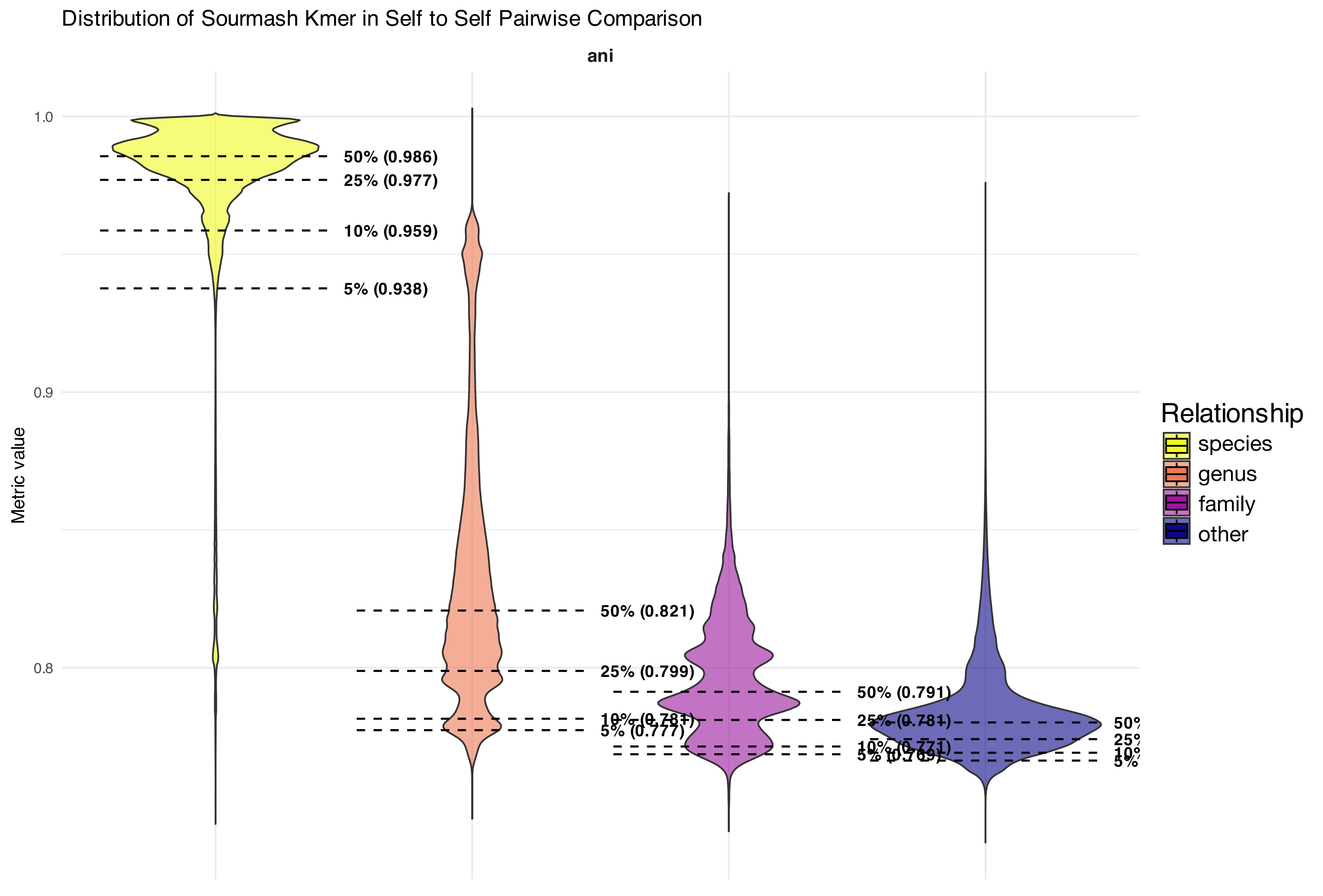

### Supplemental Figure 5

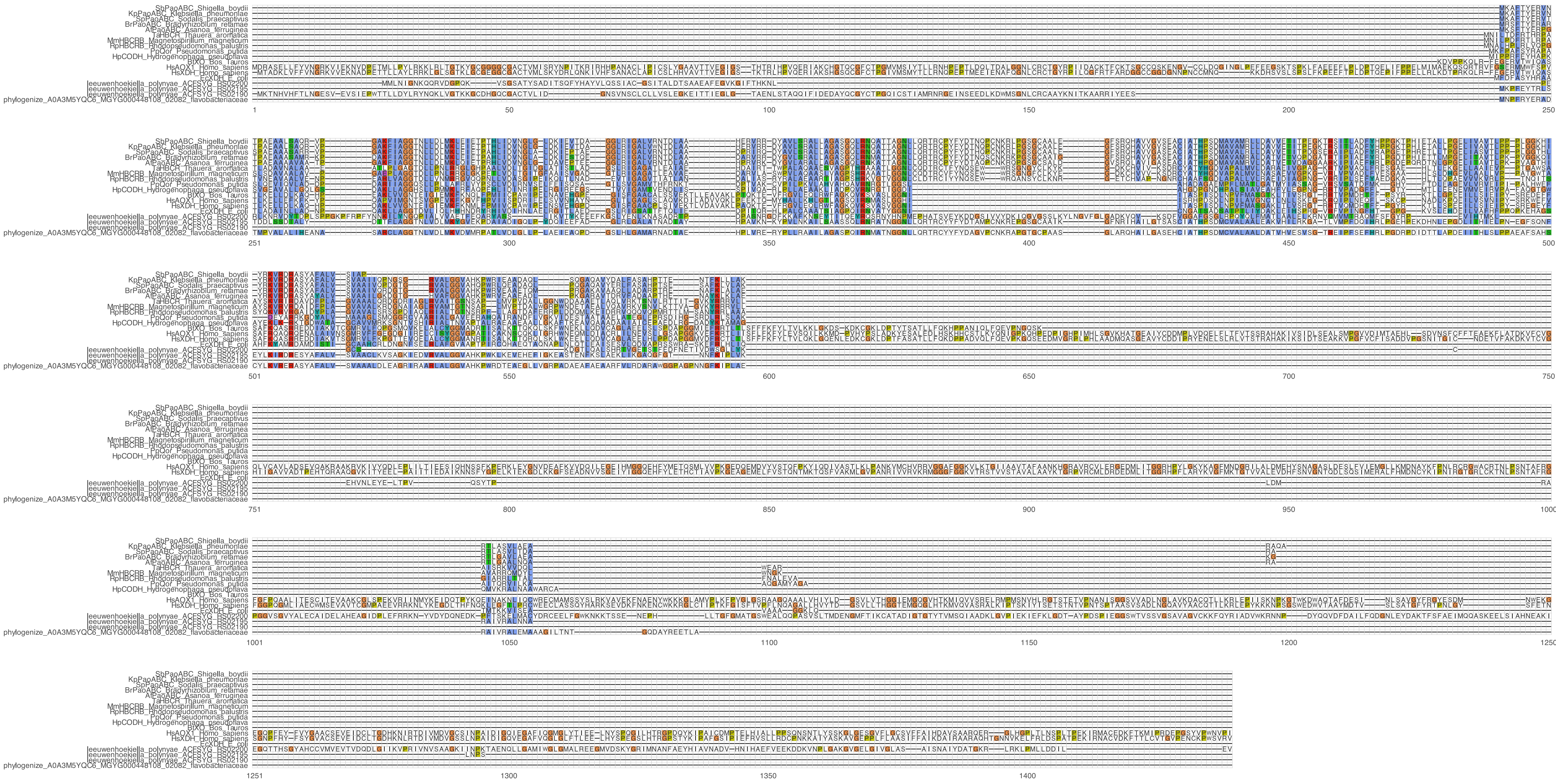
